# Empirical validation of directed functional connectivity

**DOI:** 10.1101/070979

**Authors:** Ravi D Mill, Anto Bagic, Walter Schneider, Michael W Cole

## Abstract

Mapping directions of influence in the human brain connectome represents the next phase in understanding its functional architecture. However, a host of methodological uncertainties have impeded the application of directed connectivity methods, which have primarily been validated via “ground truth” connectivity patterns embedded in simulated functional MRI (fMRI) and magneto-/electro-encephalography (MEG/EEG) datasets. Such simulations rely on many generative assumptions, and we hence utilized a different strategy involving empirical data in which a ground truth directed connectivity pattern could be anticipated with confidence. Specifically, we exploited the established “sensory reactivation” effect in episodic memory, in which retrieval of sensory information reactivates regions involved in perceiving that sensory modality. Subjects performed a paired associate task in separate fMRI and MEG sessions, in which a ground truth reversal in directed connectivity between auditory and visual sensory regions was instantiated across task conditions. This directed connectivity reversal was successfully recovered across different algorithms, including Granger causality and Bayes network (IMAGES) approaches, and across fMRI (“raw” and deconvolved) and source-modeled MEG. These results extend simulation studies of directed connectivity, and offer practical guidelines for the use of such methods in clarifying causal mechanisms of neural processing.

## 1. Introduction

The advent of network methods stands as a significant development in human cognitive neuroscience, enabling an expansion from characterizing the function of isolated brain regions to that of connections between regions and large-scale networks of regions (Craddock et al., 2013; Medaglia, Lynall & Bassett, 2015; Sporns, 2011). To date, this field of network neuroscience has been dominated by methods of “undirected” functional connectivity, which infer whether two brain regions A and B are communicating in some general fashion, as typically revealed by the Pearson’s correlation computed between their activity time series (Biswal, Yetkin, Haughton & Hyde, 1995; Friston et al., 1997). In contrast, “directed” functional connectivity (or “effective” connectivity) methods clarify asymmetries in activity flow that determine whether region A is communicating downstream to region B (connectivity A→B) or vice versa (connectivity B→A). Suggested approaches to analyzing directed connectivity in brain imaging data have included Granger causality (Roebroeck, Formisano & Goebel, 2005; Seth, 2010), directed coherence (Nolte et al., 2008), dynamic causal modeling (DCM; Friston, Harrison & Penny, 2003), conditional Bayes (Patel, Bowman & Rilling, 2006) and Bayes network methods (Mumford & Ramsey, 2014). Whilst the relative capabilities of the above algorithms to map truly “causal” or “effective” connections have been debated (Friston, 2011; Roebroeck, Formisano & Goebel, 2011), they nonetheless collectively entail a conceptual advance over undirected methods by linking cognitive operations to more precise computational mechanisms. However, widespread application of directed connectivity has been hampered by a number of methodological uncertainties, spanning the choice of imaging modality, directional algorithm, input parameters and pre-processing steps. These uncertainties call for concerted attempts to validate directed connectivity, and it is the aim of the present paper to address this need.

Prior validations have predominantly relied on the recovery of directional patterns embedded in simulated datasets. Much of this work has focused on fMRI given its present popularity, and also as it presents perhaps the clearest challenges to the application of directed connectivity. Specifically, observed BOLD signals are convolved with hemodynamic response functions (HRF), and hence offer an indirect, low-pass filtered, non-linear reflection of neuronal activity. A widely cited study by Smith and colleagues (2011) aimed to address these concerns, by using a common generative model of the fMRI BOLD response (via the biophysically plausible Buxton-Friston balloon model; Buxton, Wong & Frank,1998 Friston, Mechelli, Turner & Price, 2000) to simulate a number of fMRI directed connectivity ground truths. The recovery of these ground truths was assessed across a variety of directional algorithms. Of these, the conditional Bayes method devised by Patel and colleagues (Patel’s tau; 2006) was found to identify directed connections with the highest accuracy, albeit at an overall modest level (~65%), which raised broader questions as to the efficacy of applying directed connectivity to fMRI.

Later studies have re-examined the Smith simulations and provide a more optimistic view. For example, Ramsey and colleagues highlighted the Smith simulations’ omission of algorithms that utilize multi-subject (i.e. group-level) data to identify directed connections, as well as the suppression of likely informative non-Gaussian signal components in their forward model and pre-processing steps (Ramsey, Hanson & Glymour, 2011; Ramsey, Sanchez-Romero & Glymour, 2014). The authors hence applied their IMAGES (Independent Multiple-Sample Greedy Equivalence Search; Ramsey et al., 2010) group-level Bayes network algorithm, to the same Smith fMRI simulations, after removal of a high-pass Butterworth filter that actively suppressed non-Gaussian components in the original study, and reported a marked improvement in directionality detection accuracy (>85% Ramsey et al., 2011; Ramsey et al., 2014). Deshpande and Hu (2012) highlighted further issues in the Smith forward model, in that the original simulations failed to include an explicit delay or lag in signaling between connected regions at the neuronal level, which might have contributed to the reported poor performance of lag-based Grangercausality. Other fMRI simulations that included a realistic lag in neural signaling yielded far higher Granger detection accuracy (Roebroeck et al., 2005; Deshpande, Sathian & 2010; Wang et al., 2014). The findings of these fMRI simulations should highlight general limitations of an overreliance on synthetic approaches to directed connectivity validation, in that such simulations make generative assumptions that are open to debate, and which inevitably represent a simplification of the complexities of real imaging data.

Attempts to validate directed connectivity methods in MEG/EEG have been lacking in comparison to fMRI. This likely follows from the assumption that application of directed connectivity to MEG/EEG is fundamentally more apt, given their more direct measurement of neural activation (without complications arising from HRF convolution), sampled at a higher resolution and across a broader frequency spectrum. However, these temporal features come at the non-trivial cost of lower signal-to-noise ratio, increased non-stationarity and lower spatial resolution (i.e. the “inverse problem” of localizing neural sources for the raw sensor signal; Schoffelen & Gross, 2009). Clarifying how to address these unique challenges raises the need for directed connectivity validations in MEG/EEG as well as fMRI.

To this end, Wang and colleagues (2014) embedded directional ground truths in simulated MEG/EEG (via a neural mass model, Moran et al., 2013) and fMRI data (via the Buxton-Friston balloon model used in the Smith simulations), and compared the performance of a variety of directed connectivity algorithms across both modalities. The results support the efficacy of MEG/EEG directed connectivity analysis across a number of algorithms, as well as demonstrating comparably high detection performance in the fMRI simulations (again questioning the negative findings of the Smith simulations). Whilst these multi-modal simulations illustrate that convergent directed connectivity patterns are obtainable in MEG/EEG and fMRI data, clarification on more practical issues, such as improving signal-to-noise via preprocessing strategies and insight into significance testing are lacking. Similarly, that study failed to distinguish between sensor and source-level directed connectivity in their MEG/EEG forward model, and hence sidestepped the issue of “field spread” – the spreading of activity from a single neural source across proximal sensors, which has been shown to contaminate undirected connectivity analyses in MEG/EEG data (especially at the sensor-level; Schoffelen & Gross, 2009; Hipp et al., 2012).

Extending the undoubtedly useful fMRI and MEG/EEG simulation work calls for validations of directed connectivity in real data. Such validations are rare given the difficulty in specifying “empirical ground truth” directionality patterns in real compared to synthetic data. Whilst prior empirical directed connectivity studies have yielded interpretable results, both in fMRI (e.g. Mills-Finnerty, Hanson & Hanson, 2014; Wen, Yao, Liu & Ding, 2012; Wen, Liu, Yao & Ding,2013) and in source-modeled MEG (e.g. Astolfi et al.,2007; Cole, Bagic, Kass & Schneider, 2010; Supp et al., 2007), these reports did not seek to address antecedent questions as to the base validity of applying directional methods to brain imaging data. Of the previous empirical validations, many focus on testing one specific algorithm, such as Granger causality (Roebroeck et al., 2005), Bayes network (Ramsey et al., 2014; Plis et al., 2011), directed coherence (Gomez-Herrero et al., 2008) or DCM (Bonstrup et al., 2016), rather than the more comprehensive multi-algorithm validations undertaken by the Smith and Wang simulations. Perhaps more problematic is that few if any of these prior validations have clearly formalized an a priori ground truth directed connectivity pattern with which to evaluate performance.

The over-reliance on simulations and the limited scope of the few prior empirical validations motivated the present report, which seeks to adapt the Smith and Wang simulations to affect a multi-algorithmic, multi-modal validation of directed connectivity in real data. We collected fMRI and (anatomically constrained, source-modeled) MEG data from the same sample of subjects, as they performed the same associative memory task, involving the cued retrieval of auditory-visual stimulus pairs (Figure 1). The design of the task enabled testing of a common ground truth directed connectivity pattern that was predicated on well-established cognitive neuroscience findings and known patterns of anatomical connectivity. This capitalized on the established “sensory reactivation” effect in episodic memory research, wherein retrieval of auditory or visual information reactivates the same regions involved in perceiving those modalities (Slotnick & Schacter, 2004; Vaidya, Zhao, Desmond & Gabrieli, 2002; Wheeler, Petersen & Buckner, 2000; Wheeler et al., 2006). By manipulating whether auditory stimuli cued retrieval of visual associates (“Aud-Vis” condition) or vice versa (“Vis-Aud”condition), we sought a ground truth reversal in directed connectivity between task conditions (i.e. auditory→visual ROI connectivity in the Aud-Vis condition, and visual→auditory ROI connectivity in the Vis-Aud condition; see (Figure 2). This ground truth is substantiated by known anatomical interconnectivity between auditory and visual regions identified in animals (Cappe & Barone, 2005; Mitani et al., 1985; Schroeder & Foxe, 2005).

**Figure 1.**
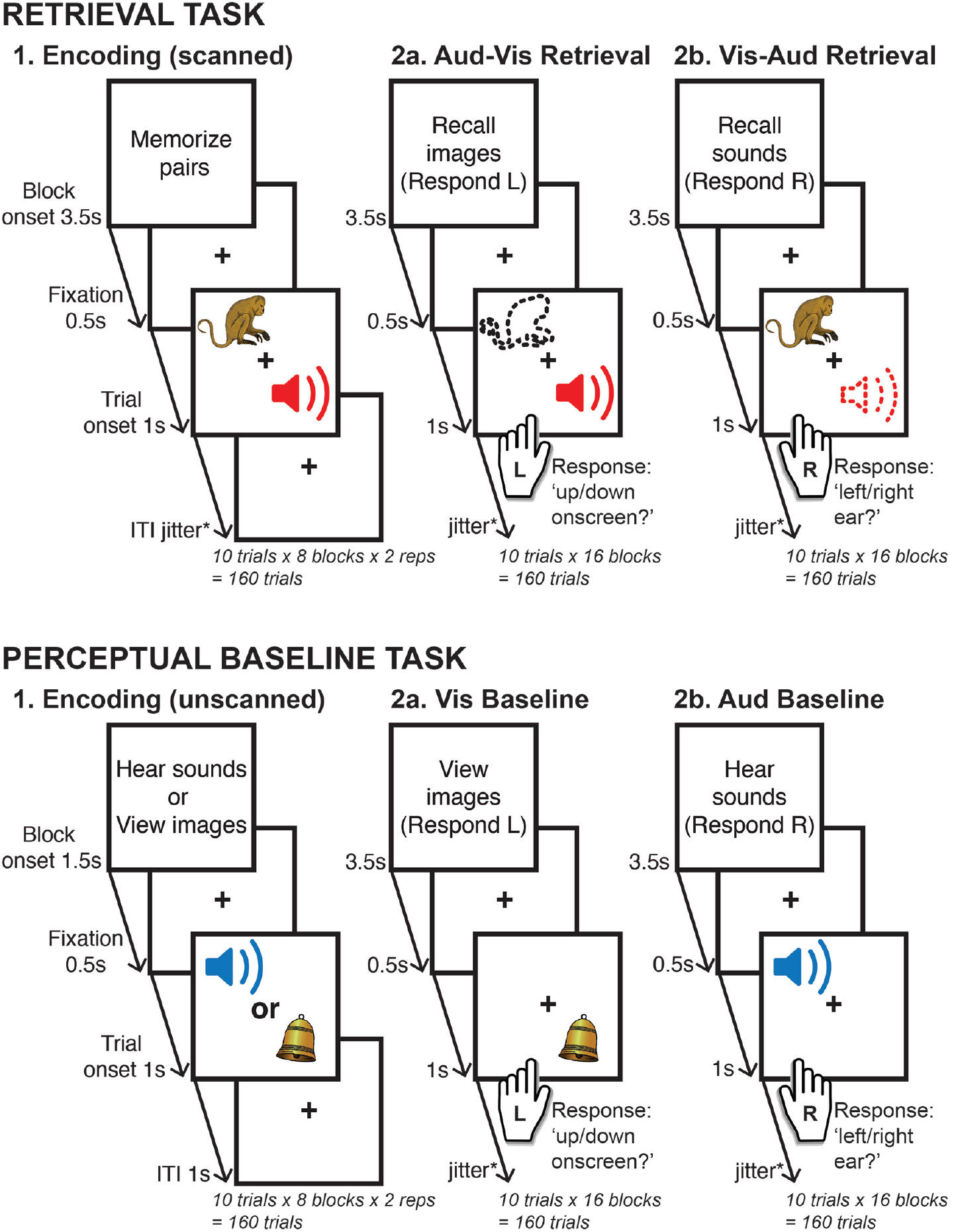
Experiment design. The retrieval task (upper panel) involved cued retrieval of auditory-visual paired associates, whereas the perceptual baseline task (lower panel) involved perception of the current auditory or visual stimulus. To clarify, the perceptual baseline task served to localize sensory ROIs for directed connectivity analyses conducted in the retrieval task. All conditions began with onset of an instruction screen that detailed the task for the ensuing block of 10 trials. Passive encoding phases were presented prior to both tasks, with the retrieval and perceptual baseline conditions randomly intermixed thereafter. The Aud-Vis retrieval task required a left-handed ‘up/down onscreen?’ decision when determining the presentation location of the episodically reactivated visual stimulus, and the Vis-Aud retrieval task required a right-handed ‘left/right ear?’ decision for the location of the reactivated auditory stimulus. This response format was maintained in the equivalent perceptual baseline conditions i.e. a left-handed decision for the presentation location of the current visual stimulus (Vis baseline) and a right-handed decision for the aural location of the auditory stimulus (Aud baseline). To enable comparisons across imaging modalities, the format of both tasks was virtually identical across fMRI and MEG scanning sessions (see Method2.2. section for details of the minor exceptions).

**Figure 2.**
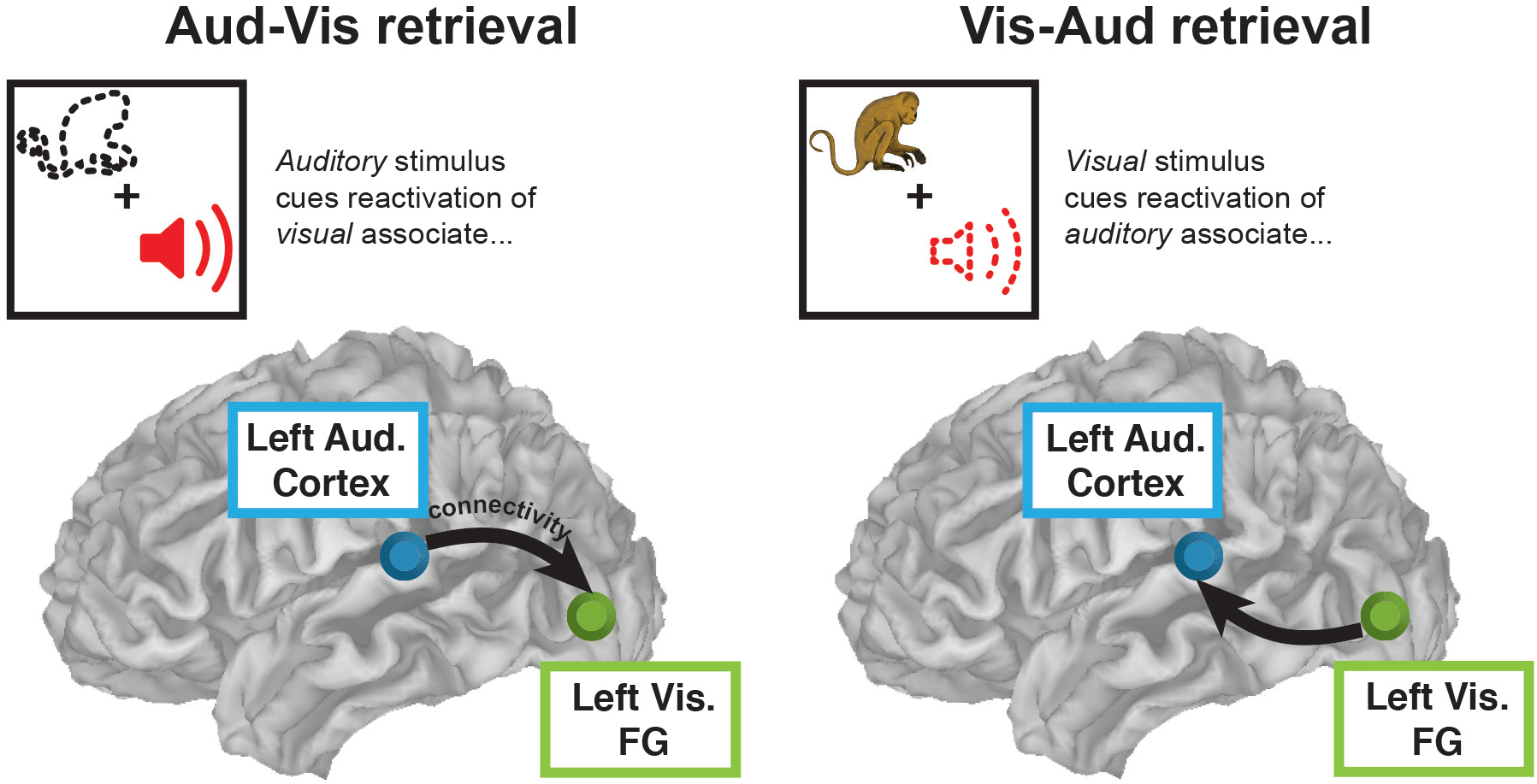
Empirical ground truth - sensory reactivation reversal. Across all tested algorithms (pairwise and multivariate) and across both imaging modalities (fMRI and MEG), this validation sought a reversal in directed connectivity between auditory and visual sensory ROIs in accordance with the reversal in task events in each retrieval condition. Arrows depict hypothesized pattern of directed connectivity. FG ‘Fusiform Gyrus’ (visual ROI).

The ground truth was tested in algorithms spanning a diverse set of assumptions, without any bias or emphasis placed on validating a given algorithm (or modality). The tested algorithms can broadly be categorized as “pairwise”, if they orient on an isolated connection-by-connection basis (Granger causality, Patel’s tau and phase slope index), or 2multivariate“, if they orient individual connections only after considering all other conditioning relationshipsamongst the region time series (IMAGES; see Supplementary Materials for descriptions of all tested algorithms)^1^).We also explored more practical issues, namely the efficacy of statistically analyzing the relative change versus absolute values of observed directed connectivity estimates (as highlighted in Roebroeck et al., 2005), the performance of fMRI algorithms withand without blind deconvolution (as highlighted in David et al., 2008), and the effect of variation in the input format and duration in the MEG analyses. The findings provide an overall positive view on the validity of applying extant directional algorithms to real human functional neuroimaging data.

## 2. Method

### 2.1. Participants

Data were analyzed for ten participants (6 female; age range = 19-46 years, mean =24.8) that underwent separate MRI and MEG scanning and had complete datasets for both, out of a total sample of 12 (two participants were excluded for problems preventing acquisition of their fMRI and MEG data respectively). Participants studied paired associates in a separate session prior to each imaging session, resulting in a total of four two-hour sessions (eight hours total per participant). Participants were recruited from the University of Pittsburgh and the surrounding area, under the exclusion criteria of any medical, neurological, or psychiatric illness, any contraindications for MRI or MEG scans, non-native English speaking or left handedness. All participants gave informed consent.

### 2.2. Design and procedure

Both fMRI and MEG experimental sessions were programmed in E-prime (Schneider et al., 2012), and comprised an identical associative memory task wherein participants encoded and then retrieved auditory-visual paired associates (see Figure 1 for design schematic). The stimuli were semantically related sounds and pictures of familiar animate and inanimate objects (e.g. horse, flute, kettle), with separate stimulus sets randomly sampled for each participant’s fMRI and MEG sessions. These sessions were conducted on separate days, with the ordering counterbalanced. Participants were only included in the main experiment if their accuracy during a practice retrieval phase (presented prior to the first encoding phase of the main experiment, see below) was greater than 80%.

Both fMRI and MEG experimental sessions comprised two main task phases - the retrieval task and the perceptual baseline task - both of which were preceded by encoding phases conducted on separate days (to facilitate stronger reactivation effects). The first encoding phase was held on the day prior to the fMRI/MEG scanning session and comprised stimuli for both retrieval and perceptual tasks. The second encoding phase for the retrieval task was scanned, whereas the second encoding phase for the perceptual task was presented prior to the scanning session (on the same day). For the retrieval task, both encoding phases involved passive viewing of simultaneously presented auditory and visual stimulus pairs. The auditory stimuli were presented to either the left or right ear, and the visual stimuli were presented on either upper or lower sections of the onscreen display. The retrieval task itself presented an auditory stimulus that cued retrieval of its encoded visual associate (the “Aud-Vis” condition), or a visual stimulus that cued retrieval of its auditory associate (the “Vis-Aud” condition). In both retrieval conditions, participants had to respond according to the source presentation location of the retrieved or “reactivated” stimulus i.e. whether the visual stimulus reactivated in the Aud-Vis condition had been presented to up/down onscreen locations at encoding, or whether the auditory stimulus reactivated in the Vis-Aud condition had been presented to left/right aural locations at encoding. Subjects made up/down onscreen decisions in the Aud-Vis condition via a response pad placed in their left hand, and left/right ear decisions in the Vis-Aud condition via a response pad placed in their right hand. Each participant’s left hand was placed perpendicular to their right hand, facilitating the up/down visual response assignments to the index and middle fingers of the left hand (and left/right auditory response assignments to the same fingers of the right hand). Participants were instructed to retrieve the pictures and sounds as vividly as possible, so as to strengthen the sensory reactivation effect.

The perceptual baseline task was matched to the retrieval task in all aspects except that responses were driven by *perception* of the cue stimulus, rather than *retrieval/reactivation* of the cue’s paired associate (see Figure 1). Both encoding phases for the perceptual task involved passively viewing visual and auditory stimuli presented in isolation, either to up/down onscreen locations or left/right aural locations respectively. These phases served to equate prior exposure to stimuli presented in the perceptual task with those presented in the retrieval task. The perceptual baseline task itself required participants to respond according to the presentation location of individually presented visual or auditory stimuli, with up/down visual onscreen decisions being made with the left hand (“Vis Baseline” condition) and left/right auditory ear decisions being made with the right hand (“Aud Baseline” condition).

All task phases were presented as 10-trial blocks, which were preceded by instruction screens notifying participants of the upcoming task, as well as the response format where required (i.e. for the retrieval and perceptual baseline conditions). The chronology of the fMRI and MEG sessions was identical: unscanned encoding for later perceptual baseline stimuli, followed by scanned encoding for later retrieval stimuli, followed by a random intermixing of retrieval and perceptual baseline mini-blocks. This format resulted in a total of 160 trials for the two retrieval conditions (160 Aud-Vis, 160 Vis-Aud) and the two perceptual baseline conditions (160 Vis Baseline, 160 Aud Baseline), for both fMRI and MEG sessions.

To clarify, the perceptual baseline and retrieval tasks were matched for sensory input, prior exposure to the stimuli (via the encoding phases accompanying each task) and response format. Task timings were also matched across retrieval and perceptual baseline tasks, with cue stimuli appearing onscreen for 1s, followed by a jittered ITI fixation cross (fMRI jitter = 2-6s, mean 4s; MEG jitter = 1-2s, mean 1.5s), with responses collected for a window extending into the ITI (fMRI response window = 3s after cue onset; MEG response window = 2s after cue onset), followed by an inter-block rest period (fMRI inter-block = 12s, MEG inter-block = 3s). Hence, the two tasks differed only in terms of whether the task was one of retrieval or perception. The close correspondence between the perceptual and retrieval tasks enabled use of the former task to localize relevant sensory ROIs for directed connectivity analyses conducted in the latter task (as detailed in later sections).

### 2.3. MRI acquisition and preprocessing

MRI image acquisition was conducted on a 3T Siemens Trio scanner. Anatomical MP-RAGE and T2 images were collected prior to the functional EPI sequence, with the latter recording 528 images in each of 10 task blocks (TR = 1000ms, no. of slices = 19, FOV = 205mm, echo time = 30ms, flip angle = 64°, voxel dimensions = 3 x 3 x 5.3mm). The encoding phase for the retrieval task constituted the first 2 fMRI acquisition blocks, and the retrieval and perceptual baseline tasks were randomly intermixed in the remaining 8 blocks. A short TR was enabled by Siemens’ generalized auto-calibrating partially paralleled sequence (Griswold et al., 2002).

Pre-processing of the fMRI data was carried out in AFNI (Cox, 1996) and FreeSurfer (Desikan et al., 2006). Each subject’s fMRI images underwent slice timing correction, before being simultaneously aligned to the start of the first EPI sequence to correct for head motion, coregistered to the subject’s anatomical MP-RAGE, and normalized to a standard MNI template. Gaussian spatial smoothing was applied at 6mm full width at half maximum. Freesurfer segmentations of the subject’s skull-stripped anatomical MRIs were used as masks to remove non-gray matter voxels.

### 2.4. fMRI GLM localizer

Regions of interest for the fMRI and MEG directed connectivity analyses were localized via a blocked fMRI general linear model (GLM), contrasting the two perceptual baseline task conditions (Vis baseline versus Aud baseline). For each participant, boxcar block onset regressors (spanning the 50 TR duration of each 10-trial task mini-block) were convolved with the canonical HRF. Separate regressors were specified for all scanned task conditions (retrieval encoding, Aud-Vis retrieval, Vis-Aud retrieval, Vis baseline, Aud baseline) to estimate voxel-wise BOLD amplitudes from the pre-processed fMRI data.

Linear contrast images were computed at the subject level for both the visual ROI localizer (Vis baseline > Aud baseline) and the auditory ROI localizer (Aud baseline > Vis baseline). The subject-level contrast images were subjected to one-sample t-tests at the group level, against the null hypothesis of a zero contrast effect at each voxel (Worsley & Friston, 1995). The group-level contrasts were corrected for multiple comparisons at the false discovery rate-adjusted (FDR; Genovese et al., 2002) level of p < .05. Hence, a visual sensory ROI was localized by the Vis baseline > Aud baseline contrast, and an auditory sensory ROI was localized by the Aud baseline > Vis baseline contrast (see Figure 3).

**Figure 3.**
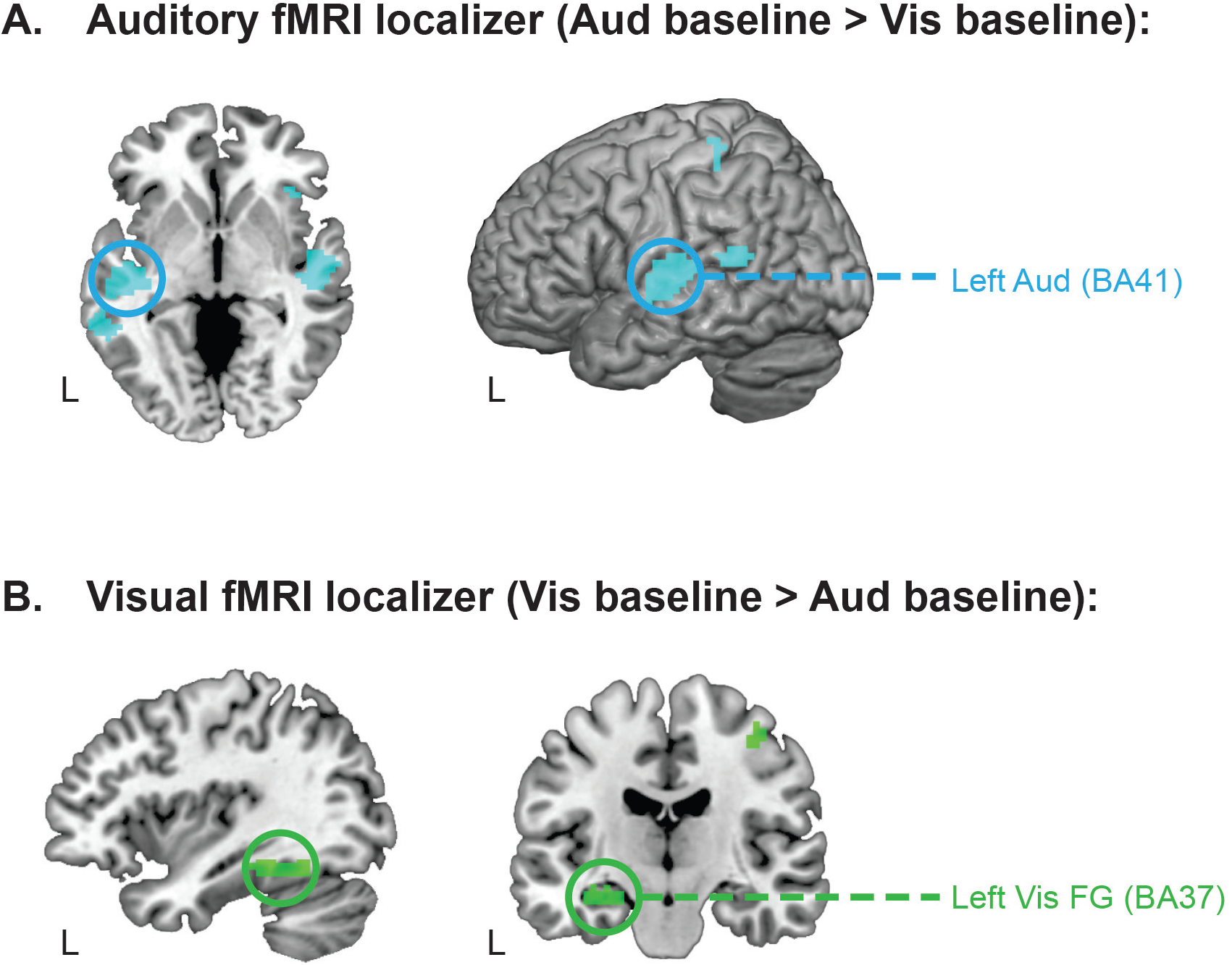
fMRI GLM localizes regions of interest for subsequent directed connectivity analyses. Panel A shows the whole-brain activation map for the perceptual baseline contrast localizing the left auditory ROI. Panel B shows the whole-brain activation map for the perceptual baseline contrast localizing the left visual fusiform (FG) ROI. All activation maps are corrected for multiple comparisons at an FDR level of p < .05 (10 contiguous voxels). ROIs from which time series were extracted for the directed connectivity analyses are circled. Left Aud = left auditory cortex; Left Vis FG = left visual fusiform gyrus; BA = Brodmann area.

### 2.5. fMRI time series extraction

fMRI time series were extracted from the ROIs localized by the GLM in identical fashion for all tested directed connectivity algorithms (detailed in the Supplementary Method). Subjects’ MNI-transformed EPI images were first converted from raw arbitrary units to a percent signal change measure, normalized to the whole-brain mean signal. Time series were extracted from each ROI as the average signal across all suprathreshold clusters at each TR. Directed connectivity was then estimated separately for Aud-Vis and Vis-Aud retrieval task conditions via the selected algorithms, on a blockwise basis for the pairwise algorithms (spanning 50 TRs in each block, excluding the block onset instruction screen and the inter-block rest period), and for the concatenated data for the multivariate IMAGES algorithm. See Table 2 for a summary of the fMRI input format and other parameters specific to each tested algorithm.

**Table 2.** Input parameters for directed connectivity algorithms applied to the “raw” fMRI data. Note that the parameters for the deconvolved fMRI analyses were identical except that Granger results are presented for a model order of 10 (see Supplementary Method for details).

| Algorithm | Estimation | Input time series | Key parameters |
| --- | --- | --- | --- |
| Granger causality (Pairwise) | Blockwise | 50 TRs x 16 blocks | Model order 2 (= 2 TRs) |
| Patel’s tau (Pairwise) | Blockwise | 50 TRs x 16 blocks | Continuous input (mapped to 0-1 range) |
| IMAGES (Multivariate) | Entire run | 800 TRs | R3 LOFS orientation ($\alpha$ threshold = .001) |

### 2.6. Deconvolved fMRI directed connectivity analysis

Directed connectivity was also analyzed in fMRI time series data that had first undergone “blind deconvolution” to remove the influence of the hemodynamic response function (HRF) from the activation time series. Significant HRF variability has been documented across brain regions and subjects (Handwerker, Ollinger & D’Esposito, 2004) and has been shown to elicit spurious estimates of directed connectivity, particularly for the lag-based Granger causality method (Deshpande & Hu, 2012; Schippers, Renken &Keysers, 2011). Methods of blind deconvolution remove HRF variability by first constructing a forward model describing how underlying neural activation leads to an observed BOLD response, as mediated by known biophysical properties of the HRF that are captured by four key parameters in the Buxton-Friston balloon model (Buxton et al., 1998; Friston et al., 2000). This forward model is then inverted to derive estimates of the four HRF parameters and the underlying neural activation that explain the most variance in the observed BOLD time series. Intuitively, this can be thought of in terms of model fitting, in which the best fitting HRF shape across all time points of a given fMRI run is used to estimate the neural activity at each individual time point. In theory, blind deconvolution yields a continuous fMRI time series that is free from HRF-induced temporal confounds.

For model inversion, we adopted the cubature kalman filtering approach described by Havlicek and colleagues (2011), as these authors demonstrated that it outperformed inversion via Bayesian filtering (used in the alternative dynamic expectation maximization method, DEM; Friston, Trujillo-Barreto, & Daunizeau, 2008) in simulated data. The raw fMRI time series were input (after demeaning) to the cubature kalman filtering algorithm, which estimated the neural activation recursively with an integration step parameter of 1/5 (as in Havlicek et al., 2011), and subjected the resulting continuous deconvolved fMRI time series to the same pairwise and multivariate algorithms as were used in analyzing the “raw” fMRI time series (see section 2.10. and Supplementary Methods for further details).

Whilst blind deconvolution has been presented primarily as a means of improving fMRI Granger causality analysis, we also conducted exploratory Patel’s tau and IMAGES analyses of the deconvolved data. As with the “raw” fMRI analyses (see Supplementary Method), Granger model order was set on both an a priori (2 TRs) and data-driven basis (10 TRs via the Bayesian Information Criterion, Schwartz, 1978). Both treatments of model order yielded comparable results, however we confine our report to deconvolved fMRI Granger analyses of the data-driven model order of 10, as these were numerically stronger. Note that the results from all variations in model order (and algorithm) were included in the FDR multiple comparisons correction procedure detailed in section 2.10.

### 2.7. MEG acquisition and preprocessing

Continuous MEG recordings were acquired on a Vectorview 306-sensor system (204 planar gradiometers, 102 magnetometers; Elekta Neuromag) across 10 task blocks (2 encoding, 8 retrieval), with a sampling rate of 1000Hz and an online low-pass filter of 330Hz. Eye movement and blink artifacts were monitored by two vertical and two horizontal electrooculogram electrodes. Electrocardiogram recordings were also made by placing two electrodes on each subject’s chest. Head position was monitored by four head position indicator (HPI) coils placed on the scalp. The three-dimensional locations of cardinal landmarks (nasion, left and right pre-auricular areas), the HPI coils and 20+ additional points on the scalp were digitized via a Polhemus tracker (Fastrak) to aid subsequent preprocessing.

Head motion artifacts were corrected with reference to the HPI coil readings of the position of each participant’s head relative to the MEG sensor taken on each run. Signal space separation (MaxFilter) was used to realign head position between runs, and to remove other external noise influences on the idealized MEG signal (Taulu, Simola & Kajola, 2005). The electrooculogram and electrocardiogram recordings were then used to regress out ocular and pulse artifacts respectively (using a 60ms sliding window; Wallstrom et al., 2004). Further MEG pre-processing was conducted within Fieldtrip (Oostenveld, Fries, Maris & Schoffelen, 2011). Trials with prominent squid jumps persisting after signal space separation were identified as high deviations in z-normalized amplitude that were common across sensors, and subsequently removed from the analysis. Line noise was removed by notch filtering, and the data was band-pass filtered into the 1-150Hz range. The trial-by-trial MEG data for each subject was finally baseline corrected relative to the 200ms preceding cue onset.

### 2.8. MEG beamformer source modeling

Source modeling of the pre-processed MEG data was conducted in Fieldtrip. Individualized head models were constructed as realistic single shells (Nolte, 2003) warped to each subject’s anatomical MRI MP-RAGE images. To capitalize on the higher spatial resolution of fMRI, MEG source locations were set as 1 cm spheres at the center of mass MNI coordinates of ROI clusters localized from the fMRI GLM^2^. Sources at these coordinates were positioned on an MNI template brain, which was warped to the subject’s anatomically constrained head model. Leadfields were computed from these source locations via Maxwell’s equations to constitute the full forward model. The linear beamformer method (LCMV; van Veen, van Drongelen, Yuchtman & Suzuki, 1997) was then used to reconstruct source activity in the time domain, in the form of an adaptive spatial filter that passes activity from source locations with unit gain whilst maximally suppressing activity from all other sources. The beamformer was calculated as a “common filter” from the sensor signal averaged over -200 to 2000ms trial epochs across both conditions of interest (i.e. Auditory-Visual retrieval and Visual-Auditory retrieval). This common filter approach ensures robust estimation of sensor covariance (as collapsing across conditions increases the number of samples) and eliminates the possibility that differences in source activation across conditions of interest are driven by differences in the underlying filters. The trial-by-trial sensor data was projected through the averaged beamformer filter to reconstruct the source time series for each ROI, for each condition. The three-dimensional dipole moment of the trial-by-trial beamformer time series was finally projected onto the x, y or z direction of maximal power via singular value decomposition.

### 2.9. MEG time series extraction

The higher sampling rate of the MEG data necessitated further processing decisions as to the *format* and *duration* of the input time series. The format of the input time series was varied as appropriate to each algorithm, between the raw signed signal (used for the spectrally resolved phase slope index method) and an unsigned, reduced root mean square power computation over 50ms time windows (50ms RMS; used for the Granger causality, Patel’s tau and IMAGES methods; see Supplementary Methods). Regarding input duration, all tested algorithms were run on two epoch lengths extracted on each trial - short epochs (0-500ms from trial onset) and long epochs (0-1000ms from trial onset). This variation in input duration was included in the correction for multiple comparisons at FDR p < .05 (see next section for further details). See Table 3 for a summary of input characteristics and other factors specific to each MEG algorithm.

**Table 3.** Input parameters for MEG directed connectivity algorithms.

| Algorithm | Estimation | Input format | Input duration | Key parameters |
| --- | --- | --- | --- | --- |
| Granger causality (Pairwise) | Blockwise | 50ms RMS | Long (0-1000ms) | Model order 2 (= 100ms) |
| Patel's tau (Pairwise) | Blockwise | 50ms RMS | Long (0-1000ms) | Continuous input (mapped to 0-1 range) |
| PSI theta (Pairwise) | Blockwise | Raw 1000Hz | Long (0-1000ms) | f segment = 250ms |
| IMAGES (Multivariate) | Entire run | 50ms RMS | Short (0-500ms) | R3 LOFS orientation ($\alpha$ threshold = .001) |

Note: whilst all algorithms were computed for long and short trial epoch durations, the parameters provided above are for the epoch duration that yielded success (Patel's tau was unsuccessful for both durations). PSI theta was computed in the 4-8Hz frequency band; 50ms RMS = input data was reduced to a root mean square power computation over non-overlapping 50ms windows; f = segmentation window for Fourier transform.

### 2.10. Overview of directed connectivity validation analyses

The same empirical “ground truth” was tested in the fMRI and MEG directed connectivity analyses, namely a reversal in directed connectivity between auditory and visual ROIs as a function of retrieval condition (i.e. auditory→visual ROI connectivity in the Aud-Vis condition, and visual→auditory ROI connectivity in the Vis-Aud condition; see Figure 2). Standard toolboxes and processing methods were used to estimate directed connectivity for all tested “pairwise” and “multivariate” algorithms. Coefficients across all pairwise algorithms were computed to enable identical interpretation of their sign. Hence, increasingly positive coefficients reflected increasing auditory → visual ROI connectivity, which should increase in the Aud-Vis compared to the Vis-Aud condition (see Figure 2). See Supplementary Methods for detailed descriptions of the tested algorithms.

Significance was interrogated as both ‘absolute’ and ‘relative changes’ in directed connectivity. In the absolute approach, the significance of directionality coefficients estimated in each *individual* retrieval condition was assessed via one-sample t-tests (against the null coefficient = 0). The relative change in directionality was assessed by *contrasting* coefficients in the Aud-Vis vs. Vis-Aud conditions via a paired-sample t-test. This relative change (or “experimental modulation”) approach has been proposed specifically as a means of controlling for artifactual influences on fMRI Granger causality (e.g. HRF variability; Roebroeck et al., 2005; Schippers et al., 2011). However, we adopt it in our analyses for all pairwise MEG and fMRI algorithms, as a general means of controlling for *any* “baseline” differences in directed connectivity between region pairs that might complicate interpretation of the individual condition estimates (e.g. due to constraints imposed by structural connectivity; Li et al., 2011). The multivariate IMAGES method does not output a quantifiable directionality coefficient, and hence we were confined to testing the absolute significance of directed graphs for each retrieval condition (as detailed in the Supplementary Methods).

In all cases, emphasis was placed on implementing a random effects approach to significance testing that would enable generalization of observed effects to the population. Hence, for the pairwise fMRI and MEG algorithms, directionality coefficients were estimated on a block-wise basis, yielding one estimate for each of the 16 mini-blocks for each retrieval condition. These block-wise condition coefficients were then averaged across blocks for each subject to enable random effects significance testing. IMAGES searches for a directed graph structure that is common to all subjects in the group, but which can vary in strength between subjects, thereby also allowing for a degree of subject variance.

All reported analyses also underwent correction for the multiple significance tests conducted across different algorithms and across any variation in algorithmic or time series input parameters. This was conducted at the FDR p < .05 level as a follow-up procedure to minimize the risk of false positives. Hence, the fMRI raw and fMRI deconvolution results were corrected across the tested algorithms and the variation in the Granger causality ‘model order’ parameter (between ‘data driven’ and ‘a priori’ selection; see Supplementary Methods). The MEG analyses underwent the same correction across algorithms and Granger model order, as well as for the variation in input duration detailed in section 2.9. (i.e. between ‘short’ and ‘long’ trial epochs). The outcomes of these FDR corrections are detailed at the end of each relevant directed connectivity result section.

## 3. Results

### 3.1. fMRI GLM localizes sensory regions of interest

We began by localizing regions of interest (ROIs) using the perceptual baseline task. We used fMRI GLMs contrasting perceptual baseline conditions, which were matched to the retrieval task in all characteristics except that they required stimuli to be evaluated based on their current perceptual location, rather than their location during encoding (see Figure.1). (Figure 3) provides the activation map for the auditory (Aud baseline > Vis baseline) and visual localizer contrasts (Vis baseline > Aud baseline), both thresholded at an FDR-adjusted level of p < .05 (with at least 10 contiguous voxels).Table 1 provides cluster details for each contrast. The left auditory cortex region identified by the auditory localizer was chosen as our auditory ROI, given prior evidence that auditory reactivation is preferentially left lateralized (Wheeler et al., 2000; Wheeler et al., 2006). The left fusiform gyrus region identified by the visual localizer has been reliably linked with visual reactivation effects in prior studies (Wheeler et al., 2000; Wheeler et al., 2006) and was hence chosen as our visual ROI. Time series were extracted from these sensory ROIs for the fMRI and MEG directed connectivity validation analyses described below.

**Table 1.** Regions active in the auditory and visual fMRI GLM localizer contrasts (both FDR-corrected at p<.05 10 contiguous voxels).

| Contrast | Region | Lat. | BA | x | y | z | Voxels |
| --- | --- | --- | --- | --- | --- | --- | --- |
| **Auditory localizer (Aud baseline > Vis baseline)** |  |  |  |  |  |  |  |
|  | Primary auditory cortex | L | 41 | -49 | -21 | 9 | 92 |
|  | Primary auditory cortex | R | 41/42 | 54 | -14 | 6 | 88 |
|  | Superior temporal gyrus | L | 22 | -61 | -44 | 14 | 34 |
|  | Precentral gyrus | L | 3 | -34 | -36 | 64 | 11 |
|  | Insula | R | 47 | 39 | 25 | 3 | 10 |
| **Visual localizer (Vis baseline > Aud baseline)** |  |  |  |  |  |  |  |
|  | Fusiform gyrus | L | 37 | -24 | -45 | 16 | 78 |
|  | Precentral gyrus | R | 3/4 | 39 | -19 | 55 | 11 |

Note: x, y and z coordinates correspond to center of mass locations for each cluster in MNI space; Lat. = lateralisation; BA = Brodmann Area.

### 3.2. Directed connectivity validation with “raw” fMRI time series

We used the sensory ROIs to empirically validate directed functional connectivity methods with fMRI. Time series were extracted from the auditory and visual ROIs and submitted to the sensory reactivation validation, which targeted a reversal in information flow between auditory and visual ROIs in accordance with manipulation of the timing of cue and reactivation events across the Aud-Vis and Vis-Aud retrieval conditions (see Figure 2). The tested fMRI algorithms spanned both pairwise (Granger causality, Patel’s tau) and multivariate (IMAGES) approaches. Table 2 details the input parameters specified for each fMRI algorithm, and Figure 4 provides the results.

**Figure 4.**
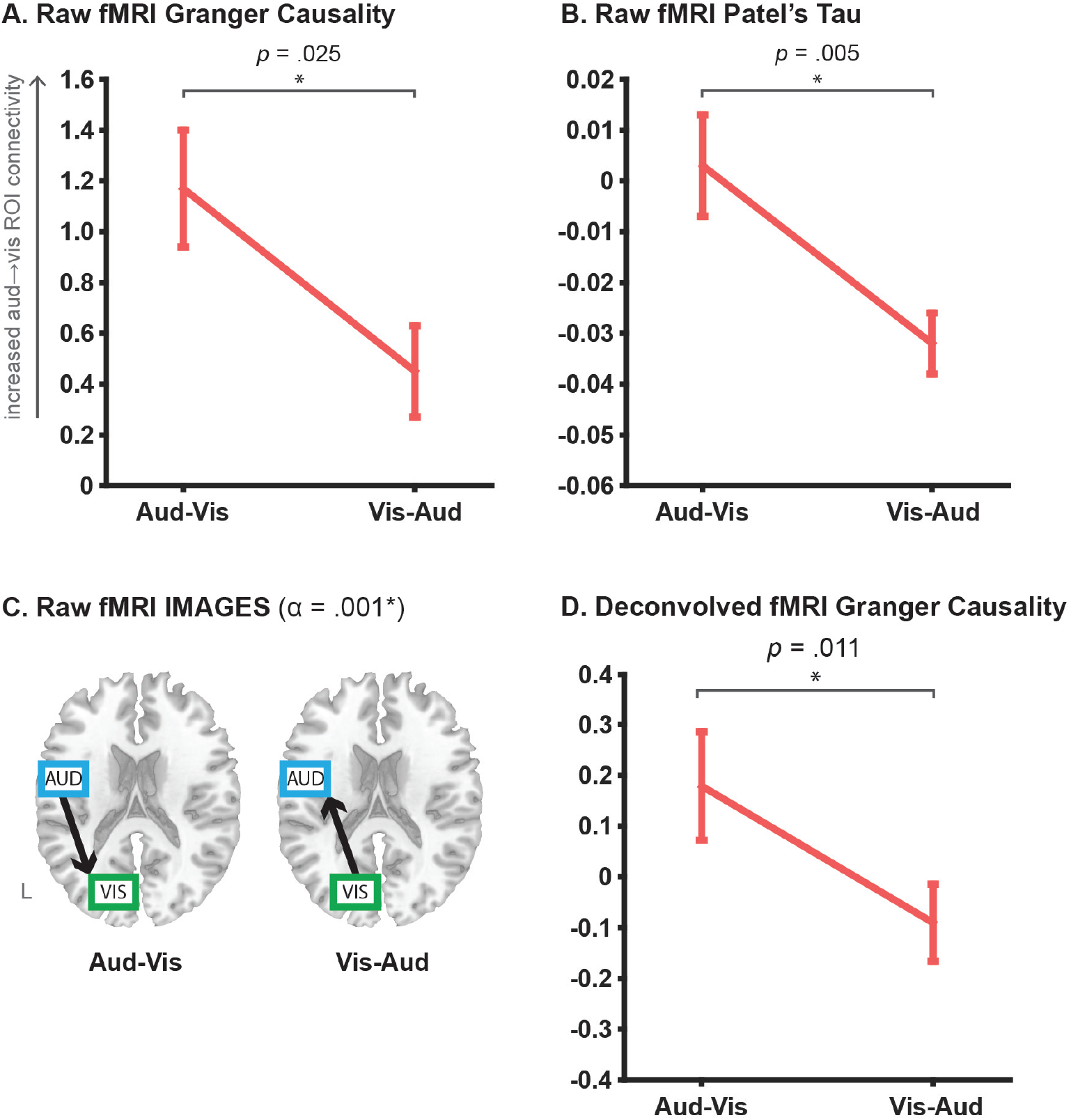
fMRI directed connectivity results. Panels A and B plot the directionality coefficients obtained for the pairwise Granger causality and Patel’s tau analyses respectively, across Aud-Vis and Vis-Aud retrieval conditions, as conducted on the “raw” fMRI signal. Error bars show standard error of mean. To clarify, increasingly positive Granger and Tau coefficients reflect increasing auditory→visual ROI connectivity, which should increase in the Aud-Vis condition. This was the pattern observed in the data across both raw pairwise methods, as revealed by the relative change in directionality (via paired sample t-test). Panel C shows the results for the multivariate IMAGES algorithm applied to the raw fMRI signal, in the form of directed graphs estimated in each individual retrieval condition, with the LOFS orientation step thresholded at an alpha level of .001. Directed graphs in both conditions were consistent with the ground truth reversal. Panel D shows the results for Granger causality applied to the deconvolved fMRI time series, with coefficients interpreted identically to the raw fMRI analyses in A and B. The deconvolved Granger analyses also yielded a significant relative change in directionality that was consistent with the ground truth. Refer to Table 2 for input parameters for all tested fMRI algorithms.

All tested fMRI algorithms recovered the ground truth directed connectivity reversal, albeit with some nuances in how this pattern was revealed. For the pairwise algorithms, the ground truth was revealed as the *relative change* in directionality between Aud-Vis and Vis-Aud retrieval conditions, rather than the *absolute* values of the coefficients estimated in each individual condition. To demonstrate via the Granger causality results in Figure 4a, whilst both condition coefficients are reliably different from 0 (via one-sample t-test, p < .05), the coefficient in the Vis-Aud condition is positive, reflecting a dominant directionality from auditory→visual ROIs. This is converse to the anticipated absolute directionality for this condition, which predicted a negative coefficient for dominant visual→auditory connectivity. Similar difficulties arise when interpreting the individual coefficients of the Patel’s tau analysis (Figure 4b). Whilst tau changed sign between retrieval conditions (more consistent with a change in the overall direction of influence between sensory ROIs), only the Vis-Aud coefficient was reliably different from 0 (at p < .05).

Rather, for both pairwise Granger and tau algorithms, contrasting the relative change in the coefficients across the two retrieval conditions yielded the correct pattern - an increase in auditoryˆvisual ROI connectivity in the Aud-Vis compared to the Vis-Aud condition. This supports prior suggestion that this relative change or “experimental modulation” approach is effective in counteracting confounding “baseline” differences in directed connectivity between pairs of regions (e.g. arising from inter-regional HRF variability; Roebroeck et al., 2005). Crucially, whilst the virtue of this approach has previously been discussed exclusively in the context of improving fMRI Granger analyses, we demonstrate here that it has a similar positive effect in improving Patel’s tau, suggesting it might be relevant to optimizing fMRI directed connectivity in a more general sense.

In contrast to the pairwise methods, the multivariate IMAGES algorithm does not output a quantifiable coefficient and hence the ground truth was interrogated as the absolute orientation of directed graphs in each condition (see Figure 4c). The algorithm’s initial search step identified an undirected connection between the auditory and visual ROIs in both retrieval conditions (final BIC scores: Aud-Vis = 213.7, Vis-Aud = 296.4). The subsequent LOFS orientation step using the R3 rule (thresholded at an alpha level of .001) recovered the ground truth directed graph in both retrieval conditions.

As detailed in the Method (section 2.10.), we also conducted a follow-up FDR correction for multiple comparisons conducted in this validation (i.e. across the tested algorithms and variation in Granger causality model order). All p values reported in the “raw” fMRI analyses survived this correction (at FDR p < .05). Overall, both the pairwise and multivariate fMRI directed connectivity algorithms were successful in recovering the sensory reactivation reversal, highlighting that fMRI can yield valid directed connectivity results.

### 3.3. Granger causality results for deconvolved fMRI time series

As detailed in the Method (section 2.6.), we also performed blind deconvolution of the raw fMRI time series to remove potentially confounding influences of HRF variability on the directed connectivity results (as has been reported previously, David et al., 2008). We used the cubature kalman filtering approach developed by Havlicek and colleagues (2011) which models subject- and region-specific HRF parameters via the Friston-Buxton balloon model (Buxton et al., 1998; Friston et al., 2000) and estimates a continuous neural activation time series that is theoretically free (i.e. deconvolved) from the HRF.

Figure 4d reveals that the deconvolved fMRI analyses also successfully recovered the ground truth reversal across retrieval conditions, as reflected by the significant relative change in Granger that was numerically higher than the equivalent raw fMRI analyses (paired t-test p = .011 for deconvolved and p = .025 for raw). Whilst the absolute coefficients in each condition were not reliably greater than 0 (via one-sample t, both p > .05), the deconvolution did at least recover the predicted directionality for each condition, which was lacking in the raw fMRI analyses (e.g. coefficient sign reflecting visual→auditory in the VisAud condition).

Whilst deconvolution has primarily been suggested as a means to improve fMRI Granger analyses (Deshpande & Hu, 2012), we also conducted exploratory analyses of Patel’s tau and IMAGES in the same data. The relative change in deconvolved Patel’s tau was in the same direction as the “raw” analyses (mean Aud-Vis = 0.021, Vis-Aud = -0.004) but failed to reach significance (paired t-test p = .112). The deconvolved IMAGES analyses correctly identified an undirected auditory-visual ROI connection in both conditions (BIC scores: Aud-Vis = 80.0, Vis- Aud = 143.5), but in both cases failed to find a reliable orientation at the LOFS stage (i.e. each graph comprised an undirected connection at alpha = .001).

As with the raw fMRI analyses, we also corrected for multiple comparisons across the tested algorithms and variation in Granger model order. The deconvolved Granger result survived this correction (at FDR p < .05). Overall, these results suggest that deconvolution is an effective means of optimizing fMRI Granger causality analysis, given that this improved the numerical strength of the significant directionality reversal, as well as corrected the sign of the individual coefficients.

### 3.4. Directed connectivity validation with MEG time series

We then tested the empirical validity of MEG directed functional connectivity methods. MEG time series were extracted from beamformer-modeled source locations set at the same MNI coordinate locations as the sensory ROIs used in the fMRI analyses. The MEG analyses targeted the same reversal in directed connectivity between sensory regions across retrieval conditions (see Figure. 2). As noted in the method however (see section 2.9.), the higher sampling rate of the MEG data led to all directed connectivity analyses being run on both long epoch (0-1000ms from trial onset) and short epoch (0-500ms from trial onset) extractions. Table 3 summarizes the input parameters for the tested MEG algorithms and Figure. 5 displays the results.

**Figure 5.**
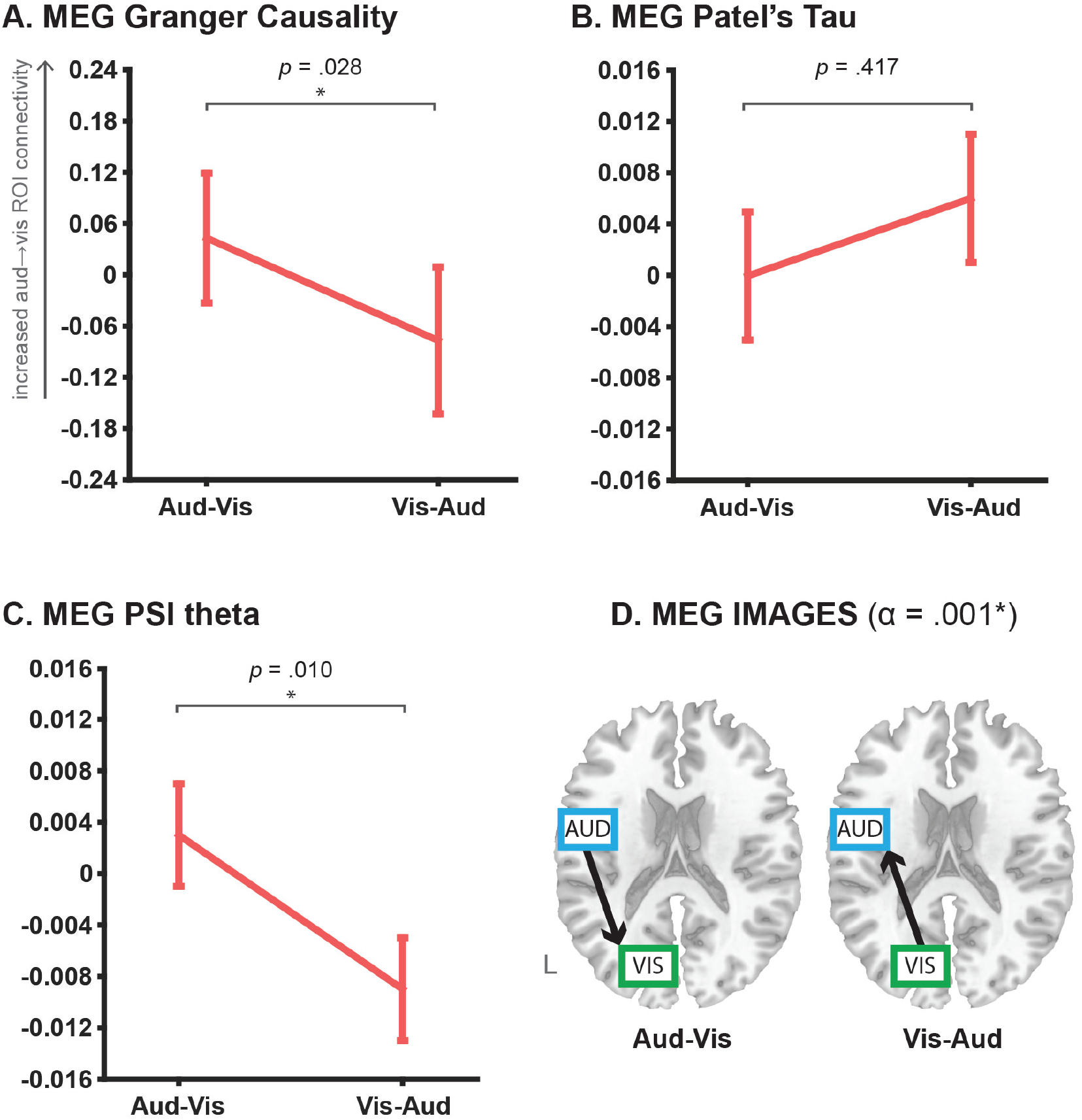
MEG directed connectivity results. Panels A, B and C plot the mean directionality coefficients obtained for the pairwise MEG Granger causality, Patel’s tau and phase slope index (PSI, confined to the theta band) analyses respectively. Error bars show standard error of mean. In panels A-C, asterisks denote significant relative changes in directed connectivity across conditions (via paired-sample t-test,<.05). For all pairwise algorithms, increasingly positive coefficients reflect increasing auditory→visual ROI connectivity, which should increase in the Aud-Vis condition. This pattern was obtained with the MEG Granger and PSI theta analyses, but not with the Patel’s tau analyses. Panel D shows the MEG results for the multivariate IMAGES algorithm, with directed graphs estimated in each individual retrieval condition at a LOFS orientation alpha level of .001. Directed graphs output for both conditions were consistent with the ground truth. See Table 3 for input parameters specified for the MEG algorithms.

The overall pattern of MEG results was comparably positive to the fMRI results, such that the ground truth reversal was obtained for two out of the three pairwise algorithms, as well as the multivariate IMAGES algorithm. Granger causality and phase slope index (PSI, confined to the theta band linked with retrieval processing; Jacobs, Hwang, Curran & Kahana, 2006; Kahana et al., 1999) both recovered the ground truth as the relative change in directionality between retrieval conditions, which reflected increased auditory → visual ROI connectivity in the Aud-Vis versus Vis-Aud conditions (see Figure 5a and Figure 5c). The reversal was reliable only for the long epoch extractions, with the short epoch extractions for the Granger coefficients (mean Aud-Vis = -0.091, Vis-Aud = 0.017; paired t-test p = .364) and PSI theta coefficients (mean Aud-Vis = 0.003, Vis-Aud = -0.010; paired t-test p = .147) both unsuccessful. As with the pairwise fMRI algorithms, the absolute individual coefficients for all MEG algorithms (except PSI in the Vis-Aud condition) failed to reliably differ from 0 at p < .05. Analyses of MEG Patel’s tau failed to obtain the ground truth directed connectivity pattern, both for the short epoch extractions (Aud-Vis = - 0.013, Vis-Aud = 0.004; paired t-test p = .09) and the long epoch extractions (displayed in Figure 5b). The results highlight that the efficacy of the relative change approach to significance testing of pairwise directed connectivity analyses extends to MEG as well as fMRI.

The MEG IMAGES analyses also recovered the ground truth pattern. This was revealed in the absolute orientations of directed graphs output for each retrieval condition, albeit only for the short epoch duration extractions (see Table 3 for input parameters). The long epoch duration analyses identified an undirected connection between sensory ROIs at the search stage for both conditions (final BIC scores: Aud-Vis = 926.9, Vis-Aud = 856.3) but then failed to orient these undirected connections via LOFS in the second stage (i.e. the final output was an undirected connection between auditory-visual ROIs in both conditions). In contrast, the short epoch duration analyses correctly identified the undirected connections (final BIC scores: Aud-Vis = 441.0, Vis-Aud = 394.5) and then oriented them in accordance with the ground truth (see Figure 5d).

We again conducted a follow-up correction for multiple comparisons in the MEG validation, across the tested algorithms, variation in Granger model order, and variation in the input duration. The PSI theta and IMAGES results survived this correction (at FDR p < .05), whereas the Granger result was marginally non-significant (FDR-corrected p = .092). Overall, the results demonstrate that convergent patterns of directed connectivity can be elucidated across different directional algorithms, as well as across fMRI and MEG neuroimaging modalities.

## 4. Discussion

The field of functional connectivity research continues to grow, encompassing recent technological innovations and multi-consortium initiatives that permit mapping of larger networks in larger samples of subjects (Hale et al., 2010; Van Essen et al., 2013). However, approaches to network analysis remain largely confined to the identification of “undirected” connections, despite the fact that alternative “directed” methods are capable of richer insight into causal mechanisms. This neglect has arisen from uncertainty over the base validity and best practices in applying directional algorithms to human imaging data, which remains despite prior efforts to validate directed connectivity in simulated fMRI and MEG/EEG (Ramsey et al., 2011; Roebroeck et al., 2005; Smith et al., 2011; Wang et al., 2014). The present report aimed to clarify some of these uncertainties by developing a principled approach to validating different directional algorithms in empirical data collected from the same subjects across different imaging modalities.

Our approach hinged on a clearly explicated ground truth, predicated on replicable cognitive neuroscience findings and established patterns of structural connectivity, which were used to evaluate the performance of algorithms and modalities. We exploited the episodic sensory reactivation effect (Slotnick & Schacter, 2004; Wheeler et al., 2000) to instantiate a ground truth reversal in auditory-visual ROI directed connectivity between task conditions (see Figure 2). This pattern was recovered across the vast majority of tested algorithms and across fMRI and MEG modalities, thereby demonstrating that convergent, empirically validated directional information can be obtained from human imaging data. These findings provide an overall more optimistic view of directed connectivity than the widely cited Smith et al fMRI simulations (2011), as well as the mixed findings from the few previous empirical validations conducted across imaging modalities (Bonstrup et al., 2016; Plis et al., 2011). These studies were also limited by their focus on validating one specific algorithm (DCM and Bayes network respectively). As such, our multi-algorithmic, multi-modal results serve as an empirical analogue of the Wang et al simulation study (2014), which reported broadly high ground truth recovery across a number of algorithms applied to synthetic fMRI and MEG/EEG data. Indeed, one of the practical guidelines emerging from our own analyses is that seeking a convergent pattern of directed connectivity across multiple directional algorithms, rather than basing interpretations solely on the findings of a single algorithm, might go some way towards validating directional results obtained in more exploratory analyses.

The present results also support the efficacy of a “relative change” or “experimental modulation” approach to significance testing in pairwise directional algorithms. Ground truth recovery was achieved by contrasting estimated coefficients between task conditions, with the absolute individual coefficient values being uninterpretable in some cases (e.g., “raw” fMRI Granger causality). Previous studies have endorsed this relative change approach in the context of controlling for the confounding influence of HRF variability in fMRI Granger causality analyses (Roebroeck et al., 2005; Roebroeck et al., 2011; Deshpande & Hu, 2012). The present findings show that the utility of this approach extends beyond this isolated context, given that it facilitated recovery of the ground truth across all tested pairwise fMRI algorithms, as well as in the pairwise MEG analyses. This suggests a role for directionality contrasts in counteracting the influence of *general* “baseline” differences in regional temporal morphologies that can confound directionality estimation. Such confounds might arise from onset or latency differences present at the neural level (Deshpande et al., 2010; Lee, Joshua, Medina & Lisberger, 2016), constraints imposed by structural connectivity (Li et al., 2011), and from any number of sources of inter-subject variability (Buxton, 2009). The importance of relative change significance testing hence stands as a key practical recommendation for others seeking to make valid inferences from pairwise directed connectivity analyses of human imaging data.

We also examined the effect of blind deconvolution on fMRI directed connectivity analysis. Modeling the underlying neural activation from the raw BOLD signal (and removing HRF variability in the process, Havlicek et al., 2011) combined with relative change significance testing to improve the numerical strength of the Granger causality results and also correct the absolute sign of the individual condition coefficients. This supports the findings of David and colleagues (2008), who only recovered a ground truth increase in directed connectivity from somatomotor to downstream regions (at seizure onset in a rat epilepsy model) via Granger causality applied to deconvolved rather than “raw” fMRI time series. Whilst we were constrained to a small sample size by the multi-modal and multi-session nature of our study, future research might clarify whether deconvolution applied to larger samples yields significant absolute coefficients, and whether newer multiband fMRI sequences (enabling sub-second TR acquisition; Feinberg et al., 2010) might also improve results. Supporting the latter point, Deshpande and colleagues (2010) demonstrated via simulations that increasing the fMRI sampling rate to approach the level of the underlying neural latency significantly improved the accuracy of Granger estimation. Nevertheless, the current findings suggest that the combination of blind deconvolution *and* a relative change approach to significance testing likely stands as the most robust strategy to fMRI Granger estimation.

It is also worth emphasizing the overall positive performance of the MEG algorithms, which except for MEG Patel’s tau recovered the ground truth. This outcome was likely aided by the rigorous source modeling procedures adopted, with the application of spatially precise linear beamforming (shown to help attenuate ‘field spread’ in MEG/EEG connectivity analyses; Schoffelen & Gross, 2009; Van Veen et al., 1997) and the use of individualized head models warped to subject’s anatomical MRIs (shown to improve source localization accuracy over templates; Klamer et al., 2015). However, it is also worth highlighting the non-trivial issue of selecting appropriate formats and durations for MEG time series input for directionality analysis, which arises collaterally from its improved temporal resolution. It was only via a principled approach at data reduction (i.e. conversion to root mean square power where appropriate) and exploring the effect of short versus long trial epoch extractions (in combination with an FDR correction procedure to minimize the risk of false positives) that MEG analyses were optimized. Whilst this principled approach yielded positive results in the present report, future studies might seek to establish a more data-driven basis for selecting optimal extraction formats and durations.

Overall, Granger causality and IMAGES could be considered the “best-performing” algorithms, given that both recovered the ground truth across fMRI and MEG modalities. As such, the positive Granger causality results accord with the Wang simulations (2014), which observed comparably high performance in directionality detection across both synthetic fMRI and MEG/EEG Granger analyses. However, more practical questions persist as to the specification of the key model order parameter. Our approach to analyze both ‘a priori’ and ‘data-driven’ means of selecting model might be a practical alternative, and it is somewhat reassuring that this variation in model order impacted on the magnitude rather than direction of the Granger coefficients.

The high performance of the Bayes network IMAGES algorithm across imaging modalities is also noteworthy. This multivariate method has not been applied to high temporal resolution MEG/EEG data (simulated or real) due to potentially serious violations to the algorithm’s underlying “i.i.d.” assumption (independent identically distributed variables; Ramsey et al., 2011 Ramsey et al., 2014) in higher temporal resolution data. We addressed this via our data reduction procedure (conversion to root mean square power over 50ms intervals), and this stands as a further practical recommendation for future analyses of Bayes network methods in MEG/EEG^3^. However, a limitation of IMAGES is the lack of a quantifiable directionality coefficient, which might prove problematic when probing relationships between directed connectivity and individual differences in behavior or clinical etiology (Craddock et al., 2013; although see Gates and Molenaar, 2012). Nevertheless, future studies might aim to extend our bivariate validations to interrogate the multivariate capabilities of IMAGES in the recovery of multi-region ground truths.

The two pairwise algorithms variously touted as “optimal” for fMRI and MEG/EEG directed connectivity analyses respectively, i.e. Patel’s tau (which performed best in the original Smith et al simulations, 2011) and PSI (which actively corrects for field spread, Nolte et al., 2008), also performed well. The MEG PSI analyses are particularly noteworthy in that these recovered band-limited directed connectivity, with reactivation effects resolved to the theta band hinting at potential “multiplexing” of directional information (Engel, Gerloff, Hilgetag & Nolte, 2013). Future validations might hence explore empirical ground truths that predict clearer segregation in directed connectivity patterns between frequency bands, as well as utilize the higher temporal resolution of MEG to examine truly dynamic or time-varying directed connectivity.

Detection of directed functional connectivity requires accurate spatial as well as temporal estimation of underlying neuronal signals. Together the fMRI and MEG results help to validate one another in this respect. This is due to the distinct set of biases underlying the spatiotemporal nature of the signals recorded by each of these imaging modalities. For instance, fMRI provides high certainty regarding the spatial location of recorded signals, yet its reliance on hemodynamic responses (which are only indirectly related to neuronal activity, and which are variable across regions and subjects) reduces temporal certainty. In contrast, MEG has low spatial certainty (e.g. due to field spread), but high temporal certainty given that it directly records neuronal electromagnetic signals. The similar results across these two modalities thus strengthen the conclusion that each provides sufficiently accurate spatiotemporal information to detect directed functional connectivity changes.

Given that both imaging modalities were successful, one might wonder whether both are necessary for directed connectivity research moving forward. We expect that future experiments (and/or simulations) will provide boundary cases for use of each modality. For instance, it may be that fMRI is unable to detect directed connectivity changes that occur dynamically within a trial, such as a reversal between the first and second hundred milliseconds of processing a visual stimulus (Pascual-Leone & Walsh, 2001). In contrast, MEG’s high temporal resolution may provide evidence for such an effect. However, directed connectivity between two neighboring occipital visual regions may be difficult to detect with MEG due to spatial proximity of the relevant signals, but this may be detectable with fMRI. These examples illustrate the need for both modalities moving forward, as well as the need for new non- invasive technologies to allow detection of directed connectivity effects that both modalities are likely blind to (e.g. changes in directed connectivity between neighboring cortical columns with 10 ms inter-signal lag).

## 5. Conclusion

Beyond clarifying the base validity of directed connectivity analysis across algorithms and imaging modalities, our findings raise certain practical guidelines for future empirical work. Firstly, analysis of the relative change in directionality coefficients across task conditions (rather than the absolute values of individual coefficients) has an overarching positive effect in controlling for baseline time series confounds, and serves to optimize directionality estimation across algorithms and modalities. Secondly, deconvolution is effective in improving fMRI Granger causality results, albeit only in combination with relative change significance testing (although larger samples might yield more interpretable absolute coefficients). Thirdly, anatomically constrained, source-modeled MEG raises the potential for spatiotemporally precise directed connectivity estimation, although care should be taken in specifying the format and duration of the high resolution input time series. Finally, the Granger causality and IMAGES algorithms performed “best” across the two imaging modalities, and seeking convergent results across directional algorithms spanning different assumptions might serve to validate more exploratory analyses.

Overall, the present findings should highlight that methodological validations in real data stand as an important precursor to functional extensions of directed connectivity, such as mapping directionality amongst multiple regions spanning the whole brain, analyzing spontaneous directionality patterns in the resting state, and interrogating relationships with individual differences. The functional refinement afforded by such validations is made clear by the present results, which directly link the reactivation of sensory information from memory to changes in directed connectivity. Further validations and functional applications of directed connectivity methods will likely furnish even greater insight into causal mechanisms of neural computation.

## Acknowledgements

We would like to thank Stephen Hanson, Catherine Hanson, Dana Mastrovito, Andreea Boston, Mark Wheeler, and Robert Kass for helpful feedback. M.W.C. was supported, in part, by a National Science Foundation IGERT grant, as well as National Institutes of Health K99-R00 grant MH096801. The content is solely the responsibility of the authors and does not necessarily represent the official views of the National Science Foundation or National Institutes of Health.

## Footnotes

1 Note that, given recent emphasis on large network graphs for characterizing brain “connectomes” (Sporns, 2014), we focus on directional algorithms that are not limited to modeling a small number of regions (unlike, e.g., DCM).

2 Directed connectivity analyses were also run on MEG time series extracted from sources set at the peak MNI coordinates of the fMRI GLM clusters, which yielded an identical pattern of results.

3 One could also speculate that the 3-cycle clique elimination constraint in IMAGES served to further counteract the field spread problem in MEG connectivity analyses, which might account for the better performance of MEG IMAGES compared to MEG Patel’s tau (despite both algorithms sharing Bayesian features). However, the algorithms also differ in that IMAGES uses Gaussian signal components to identify the core undirected graph (Step 1) and non-Gaussian components to orient this graph (Step 2 LOFS), whereas Patel’s tau uses only Gaussian information for orientation. The better performance of IMAGES compared to tau in the MEG data might therefore be consistent with prior suggestion that directional information is carried preferentially by non-Gaussian information (Mumford & Ramsey, 2014).

## Notes

**Conflict of Interest:** None

